# Normative values of neuromelanin-sensitive MRI signal in older adults obtained using a standard protocol for acquisition and analysis

**DOI:** 10.1101/2022.06.02.493125

**Authors:** Rami Al Haddad, Mira Chamoun, Christine L Tardif, Synthia Guimond, Guillermo Horga, Pedro Rosa-Neto, Clifford M Cassidy

## Abstract

**Background:** The integrity and function of catecholamine neurotransmitter systems can be assessed using MRI sequences often referred to as neuromelanin-sensitive MRI (NM-MRI). The relevance of this method to neurodegenerative and psychiatric disorders is becoming increasingly evident, and it has potential as a clinical biomarker. To support such future applications, we report here the normative range of NM-MRI signal and volume metrics in cognitively normal older adults.

**Methods:** 3 Tesla NM-MRI images and demographic and cognitive data were available from 152 cognitively normal older adults aged 53-86 years old at baseline; a subsample of 68 participants also had follow-up NM-MRI data collected around one-year later. NM-MRI images were processed to yield summary measures of volume and signal (contrast-to-noise ratio, CNR) for the substantia nigra (SN) and locus coeruleus (LC) using a recently developed software employing a fully automated algorithm. The extent of annual change in these metrics was quantified and tested for significance using 1-sample t-tests.

**Results:** Baseline SN signal (CNR) was 10.02% (left SN) and 10.28% (right) and baseline LC signal was 24.71% (left) and 20.42% (right). The only NM-MRI metric to show a significant annual change was a decrease in left SN volume.

**Conclusion:** We report normative values for NM-MRI signal and volume in the SN and LC of cognitively normal older adults and normative values for their change over time. These values may help future efforts to use NM-MRI as a clinical biomarker for adults in this age range by facilitating identification of patients with extreme NM-MRI values.

## Introduction

Neuromelanin-sensitive MRI (NM-MRI) is an increasingly popular method to interrogate the integrity and function of the brain’s major catecholamine nuclei, the dopaminergic substantia nigra (SN) and the noradrenergic locus coeruleus (LC).^1–5^ The method has been most broadly applied in research studies of neurodegenerative illnesses such as Parkinson’s disease and Alzheimer’s disease as a marker of degeneration of these nuclei.^2,3,6,7^ More recently it has been applied in psychiatric conditions, where the NM-MRI signal has been shown to correlate to important aspects of psychopathology, consistent with evidence that the signal may assay aspects of the function of these neurotransmitter systems, even in the absence of neurodegeneration.^1,8,9^

Many research studies support the potential of NM-MRI as a neuroimaging biomarker that could prove useful in clinical applications such as diagnosis or treatment of neurodegenerative and psychiatric disorders.^2,8,10^ Supporting such applications, NM-MRI has excellent test-retest reliability with Intraclass Correlation Coefficient (ICC) values of 0.95 or higher, suggesting that quantitative measures of NM-MRI signal can be consistently obtained with minimal error.^1,11,12^ Nevertheless, there are also challenges that must be overcome to support its use as a biomarker.^8,13,14^ First of all, to be used universally, it is necessary for NM-MRI metrics from different scanners and different sites to be comparable.^8,12,15,16^ This is challenging because, in many applications, it is the signal contrast of the nuclei that would be the principal metric of interest.^11,16,17^ Compared to, for instance, volumetric measures, signal contrast is even more sensitive to factors such as scanner model, pulse sequence, acquisition parameters, and head coil.^8,12^ Thus, these sources of variability should be minimized and also data harmonization methods such as ComBat should be applied as part of data processing in the context of multisite studies or when comparing NM-MRI measurements to an external standard.^1,18^ The second challenge is the observation that the NM-MRI signal increases with age, sharply during childhood, reaching a plateau in middle age, and then declining in later life.^19–21^ These findings may be due to NM’s tendency to gradually accumulate in the cell bodies of catecholamine neurons, only being cleared from the tissue after cell death.^3,19,22,23^ Indeed, post-mortem observations have found a similar pattern of age-related change in the accumulation of NM in these nuclei.^24–26^ As a result, a signal contrast that is relatively high for the SN of a teenager would be relatively low for a middle-aged adult.^19–21,27^ Therefore, age is a critical factor that should be adjusted for when using NM-MRI.^19,28^ With these challenges in mind, future NM-MRI work using standard methods of acquisition and analysis and stratifying a large sample of participants by age would be well positioned to determine the normative range of SN and LC contrast ratios and volumes in healthy people of all ages. In the context of its potential use as a biomarker, access to a standardized, age-adjusted normative range of NM-MRI metrics is essential to identify patients on the extreme ends of the range, presumably those who may require clinical intervention.

To advance NM-MRI towards its potential use as a biomarker, we conducted a NM-MRI study in a large sample of cognitively normal older adults using standard methods for acquisition and analysis via a fully automated software for processing NM-MRI images that was recently developed. While a sample larger than the present one and spanning the entire lifespan will be ultimately necessary to provide granular age adjustment, in the current study, we propose that this sample will be adequate to approximate the normative range of NM-MRI metrics in cognitively normal older adults over 53 years old, a clinically relevant age-range given the emergence of common neurodegenerative conditions in older adults. We report measures of NM-MRI signal and volume for the SN and LC, as well as their annual change.

## Methods

### Participants

Study participants from the community were enrolled in the Translational Biomarkers of Aging and Dementia (TRIAD) cohort, McGill University Research Centre for Studies in Aging, Montreal, Canada.^29^ The participants had a detailed clinical assessment, including the Clinical Dementia Rating Scale (CDR) and Mini-Mental State Examination (MMSE). All participants in the current study were deemed to be cognitively unimpaired, with no objective cognitive impairment and a CDR score of 0. Participants were excluded if they had other inadequately treated conditions, active substance abuse, recent head trauma, or major surgery, or if they had MRI/PET safety contraindication. This study was approved by the Douglas Mental Health University Institute Research Ethics Board and written informed consent was obtained from all participants.

### MRI Acquisition

All neuroimaging data were acquired at the McConnell Brain Imaging Centre of the Montreal Neurological Institute (MNI). MRI was acquired on a 3T Prisma-Fit scanner. NM-MRI images were collected via a turbo spin echo (TSE) sequence with the following parameters: repetition time (TR)=600 ms; echo time (TE)=10 ms; flip angle=120°; turbo factor=4; in-plane resolution=0.6875×0.6875 mm^2^; partial brain coverage overlaying the pons and midbrain with field of view (FoV)=165×220; number of slices=20; slice thickness=1.8 mm; number of averages=7; acquisition time=8.45 min. The image stack was aligned perpendicular to the axis of the brainstem in the region of the pons. Whole-brain, T1-weighted MR images (resolution=1 mm, isotropic) were acquired using an MPRAGE sequence for preprocessing of the NM-MRI and PET data. Quality of MRI images was visually inspected for artifacts immediately upon acquisition. In case of suboptimal image quality, scans were repeated, time permitting.

### Preprocessing of NM-MRI images

Preprocessing of NM-MRI images was performed using a fully-automated, cloud-based software package, NM-101, version 1.0.3 (Terran Biosciences). The initial algorithm steps were necessary for processing of both LC and SN metrics. These included brain extraction of T1-weighted images, spatial normalization of T1-images into standardized MNI space, rigid coregistration of NM-MRI images to T1-weighted images, and spatial normalization of NM-MRI images into MNI space (resampled at 1 mm, isotropic).

To calculate SN metrics from the spatially normalized images, we used an approach similar to our prior work; see Wengler et al., for more detailed description.^1,11,12,30^ Note that here we refer to the SN, but our SN mask also contains the dopaminergic ventral tegmental area (VTA), and thus the structure assayed here could also be referred to as the SN-VTA complex. Intensity normalization determined contrast-to-noise ratio (CNR) for each subject and voxel *v* as the relative change in NM-MRI signal intensity I from a reference region RR of white matter tracts known to have minimal NM content, the crus cerebri. Images were then spatially smoothed with a 1-mm full-width-at-half-maximum Gaussian kernel. Finally, an overinclusive mask of the SN in MNI space was applied that ensures inclusion of relevant nuclei for all individuals. SN signal was calculated by averaging CNR values for all SN voxels on the left and right sides of this mask. SN volume on the left and right sides was calculated by counting the number of SN voxels in MNI space above a fixed intensity threshold and multiplying by the volume of one voxel (1 mm^3^). The intensity threshold was equal to two times the standard deviation calculated from all reference region voxels of all subjects after trimming reference region voxels with extreme intensity values (this approach was adapted from our previous work).^12^

To calculate LC metrics from the raw NM-MRI images, we used an approach similar to our prior work; see Cassidy et al., 2022 for more detailed description.^31^ An LC search space (referred to as the LC search mask, spanning 5 mm in the rostrocaudal axis from MNI space) was created to cover the middle LC (where signal intensity is highest and automated segmentation accuracy is very high, ∼99% accuracy) and exclude the rostral and caudal portions of the structure. This LC search mask was then warped to NM-MRI space using the inverse transformations generated in the spatial normalization and rigid coregistration steps. This warped LC search mask defined a search space wherein to find the LC for each participant. A cluster-forming algorithm was used to segment the LC within this space. This was repeated on each side (left and right) and axial slice. To minimize partial volume effects, only the brightest of these voxels (peak intensity voxel) was retained for calculation of LC signal on each side and slice (on the assumption that the peak intensity voxel had the highest fraction of LC tissue. CNR for each voxel *v* in a given axial slice was then calculated as the relative difference in NM-MRI signal intensity *I* from the portion of a reference region in the same axial slice. We used a reference region with low NM concentration, the central pons. The central pons mask was transformed from MNI space and transformed to NM-MRI space for each participant. LC signal was calculated by averaging CNR values from the all peak intensity LC voxels on the left and right sides.

To calculate LC volume, the number of voxels above a fixed intensity threshold were counted on one side for every axial slice, then divided by the number of axial slices to estimate average LC voxels per slice, and then multiplied by the area per voxel. This area measure was multiplied by a fixed LC length (equal to the length of the LC search mask in MNI space, 5 mm). Therefore, all variability in our measure of LC volume was due to variability in LC area, with rostrocaudal length fixed due to the inability of our method to precisely measure this dimension. This was repeated on both the left and right sides. This method for LC volume calculation was designed to capture volume loss in neurodegenerative illness, consistent with the intent of the NM-101 software. The intensity threshold was set as the maximum value where no cognitively normal participants had floor values (0 voxels) for LC volume. Many participants had ceiling values for LC volume using these thresholds. Due to left-right asymmetry in the NM-MRI images, different thresholds were applied on the left and right sides.

### Statistical Analysis

All NM-MRI metrics generated by NM-101 software were analyzed using Matlab software. Normality was assessed using Lilliefors test (alpha=0.05). Linear regression was used to relate baseline and change NM-MRI metrics to age. Annual change metrics for all NM-MRI metrics were calculated by subtracting NM-MRI metrics at baseline from the same metrics at follow-up, 9-16 months later. 1-sample t-tests were conducted to determine if these change metrics significantly differed from 0.

## Results

### Section 1. Normative metrics: baseline

The sample included 152 adults aged between 53 and 86 years at baseline. Mean age was 71.2±5.9 years, mean years of education was 15.3±35, mean score on the mini-mental exam was 29.2±1.4, 50 participants were male (32.9%), and 110 participants were Caucasian (86.6%). Baseline SN signal had CNR values of 10.02% (left SN) and 10.28% (right SN) and baseline LC signal was 24.71% (left) and 20.42% (right; see Table 1 for all signal and volume measures). All signal measures were normally distributed at baseline (Lilliefors test, alpha>0.05). This was not the case for all volume measures (consistent with our decision to set volume thresholds where many healthy individuals would be at-or-near the ceiling, this was particularly true of LC volume metrics). Refer to Table S1 for values of all baseline summary metrics for all participants. Figure 1 shows an SN map illustrating voxelwise values for baseline SN signal.

**Table 1.** Summary NM-MRI metrics at baseline and change over 1 year

|  |  | SN |  |  | LC |  |  |
| --- | --- | --- | --- | --- | --- | --- | --- |
|  |  | Baseline | Annual change | T stat (annual change) | Baseline | Annual change | T stat (annual change) |
| Signal (CNR) | Left | 10.02±1.48 | -0.39±1.32 | -1.89 | 24.71±4.88 | 0.67±3.29 | 1.28 |
|  | Right | 10.28±1.51 | -0.30±1.18 | -1.61 | 20.42±4.95 | -0.62±2.92 | -1.35 |
| Volume (mm<sup>3</sup>) | Left | 576.4±83.0 | -30.44±75.9 | -2.57* | 6.31±1.22 | 0.27±0.92 | 1.83 |
|  | Right | 540±74.9 | -17.21±64.2 | -1.72 | 6.30±1.20 | 0.30±1.37 | 1.41 |

\*p=0.014

**Figure 1.**
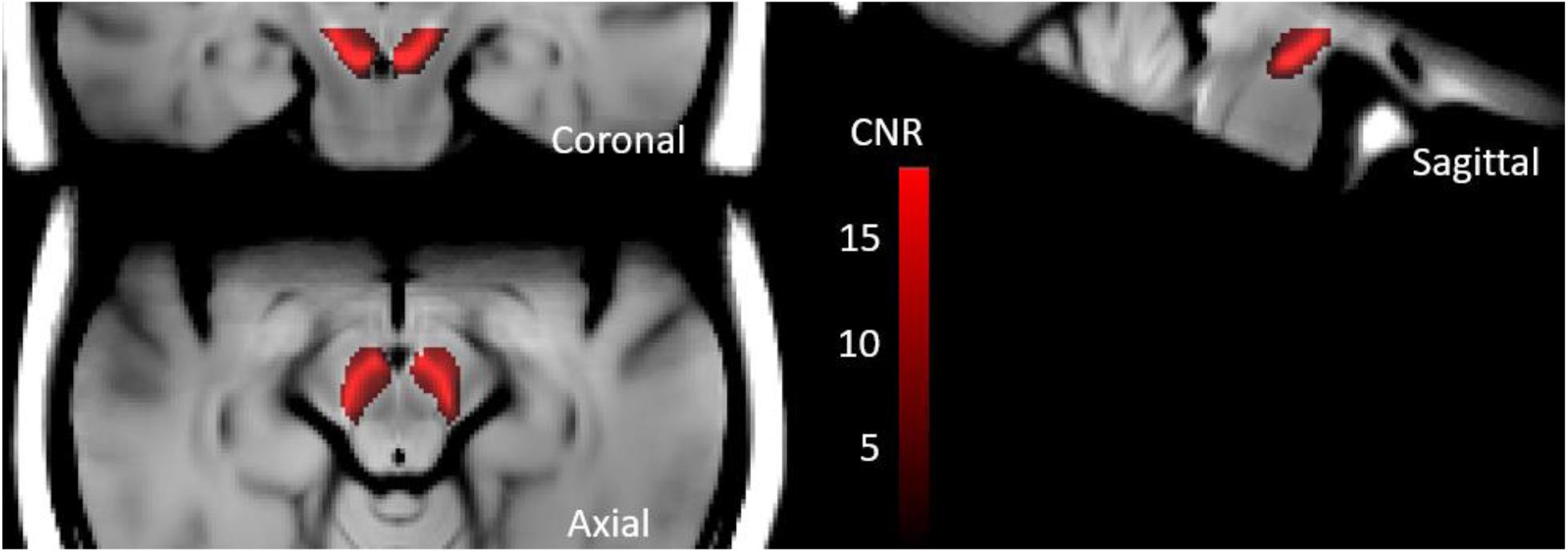
Mean NM-MRI signal from SN voxels at baseline in older adults. Signal is highest in the central SN.

Given our decision to determine normative NM-MRI metrics from a relatively broad age range, we checked whether there was meaningful variation in these metrics over this range. We confirmed that there was no significant relation of NM-MRI signal or volume in the SN or LC to age (p values ranged from 0.14 to 0.99, linear regression analysis), supporting the acceptability of this broad range and consistent with evidence that the age-related change in NM-MRI metrics plateaus in later life.^19^

### Section 2. Normative metrics: annual change

Follow up images after 1 year (9-16 months) were available for n=41 participants (n=40 for LC). Mean change in NM-MRI metrics over the year were all negative (decreases over time) for SN metrics, and tended to be positive (increases) for LC metrics (Table 1). There was a large amount of variability across subjects in all of these metrics with some subjects showing increases and some decreases over time. Similar to observations at baseline, annual change in signal metrics was normally distributed but change in volume measures was not. Only SN volume on the left side showed significant annual change across subjects (Table 1, 1-sample t-tests).

We also checked if age significantly affected annual change metrics and found this was not the case (p values ranged from 0.16 to 0.96, linear regression analysis) suggesting that, similar to baseline metrics, collapsing older adults across this wide age range may be acceptable when considering annual change metrics.

## Discussion

Here we report normative values for baseline NM-MRI signal and volume in the LC and SN and their change over time that were obtained using a fully automated algorithm. Annual change for NM-MRI summary metrics where largely non-significant but tended to decrease for SN metrics and increase for LC metrics.

These normative values from cognitively normal older individuals could be useful if NM-MRI achieves its potential as a biomarker to support clinical decision-making. For instance, NM-MRI has been widely used in research studies in Parkinson’s disease but could potentially be introduced to the clinic to assist with diagnosis or treatment monitoring by flagging individuals with low NM-MRI signal or with marked signal loss over time relative to the norms from healthy individuals. When possible, these normative values would be adjusted in a scanner-specific manner (for instance using ComBat) to account for differences across scanners that could shift the range of the NM-MRI signal, even when parameters and equipment are closely matched between scanners. The subject-level data in Table S1 could assist with this harmonization process.

Although this study was not designed to assess the performance of the algorithm, the sensitivity of the algorithm to detect SN volume changes in the expected direction over a relatively brief time interval provides support for its good performance.

The main limitation of this study is the sample size which prevented us from subdividing it into smaller age subgroups or dividing it by sex or other demographic factors. Ideally, a future study should collect a substantially larger sample of people across the entire lifespan and divide them into age subgroups (perhaps 10 years or smaller) and demographic groups. This would allow for more fine-grained norms, potentially narrowing their range and facilitating identification of patients at risk of neurodegenerative or perhaps psychiatric illness. Nevertheless, it is reassuring that we did not observe significant age-related change in any of our summary NM-MRI metrics over the age range employed here, supporting the utility of these norms for adults aged between 53 and 86 years old.

It is also important to note that these normative values are not only limited to a specific population (older adults) but also by the NM-MRI acquisition protocol. While they should be robust to small changes in sequence parameters, assuming the values are harmonized using ComBat or a similar approach, they cannot be applied universally to any NM-MRI acquisition. We report values using a TSE sequence and these values cannot be compared to values obtained using the other commonly used sequence for NM-MRI, a 2D gradient recalled echo sequence with magnetization transfer pulse (2D-GRE with MT). We selected the TSE sequence here given our objective of providing norms that could be clinically useful and the fact that on some scanner brands a developer-made 2D-GRE sequence is needed, making the latter sequence a step further removed from clinical implementation.

In conclusion, our results provide normative values of NM-MRI signal and volume for the SN and LC in cognitively normal older adults. Such norms represent one of the necessary components in the effort to introduce this promising neuroimaging method into the clinic as a biomarker for neurodegenerative and psychiatric disorders.

## References

1. Cassidy, C. M. et al. Neuromelanin-sensitive MRI as a noninvasive proxy measure of dopamine function in the human brain. Proc. Natl. Acad. Sci. 116, 5108–5117 (2019).

2. Wang, J. et al. Neuromelanin-sensitive magnetic resonance imaging features of the substantia nigra and locus coeruleus in de novo Parkinson’s disease and its phenotypes. Eur. J. Neurol. 25, 949–e73 (2018).

3. Fedorow, H. et al. Neuromelanin in human dopamine neurons: comparison with peripheral melanins and relevance to Parkinson’s disease. Prog. Neurobiol. 75, 109–124 (2005).

4. Chen, X. et al. Simultaneous imaging of locus coeruleus and substantia nigra with a quantitative neuromelanin MRI approach. Magn. Reson. Imaging 32, 1301–1306 (2014).

5. Simões, R. M. et al. A distinct neuromelanin magnetic resonance imaging pattern in parkinsonian multiple system atrophy. BMC Neurol. 20, 432 (2020).

6. Bae, Y. J. et al. Imaging the Substantia Nigra in Parkinson Disease and Other Parkinsonian Syndromes. Radiology 300, 260–278 (2021).

7. Cassidy, C. et al. Neuromelanin-Sensitive MRI as an Index of Norepinephrine System Integrity in Healthy Aging, Mild Cognitive Impairment, and Alzheimer’s Disease. Biol. Psychiatry 87, S424–S425 (2020).

8. Sulzer, D. et al. Neuromelanin detection by magnetic resonance imaging (MRI) and its promise as a biomarker for Parkinson’s disease. NPJ Park. Dis. 4, 11 (2018).

9. Paloyelis, Y., Mehta, M. A., Kuntsi, J. & Asherson, P. Functional magnetic resonance imaging in attention deficit hyperactivity disorder (ADHD): a systematic literature review. Expert Rev. Neurother. 7, 1337–1356 (2007).

10. Wang, L. et al. Substantia nigra neuromelanin magnetic resonance imaging in patients with different subtypes of Parkinson disease. J. Neural Transm. Vienna Austria 1996 128, 171–179 (2021).

11. Wengler, K., He, X., Abi-Dargham, A. & Horga, G. Reproducibility assessment of neuromelanin-sensitive magnetic resonance imaging protocols for region-of-interest and voxelwise analyses. NeuroImage 208, 116457 (2020).

12. van der Pluijm, M. et al. Reliability and Reproducibility of Neuromelanin-Sensitive Imaging of the Substantia Nigra: A Comparison of Three Different Sequences. J. Magn. Reson. Imaging 53, 712–721 (2021).

13. Trujillo, P. et al. Contrast mechanisms associated with neuromelanin-MRI. Magn. Reson. Med. 78, 1790–1800 (2017).

14. Liu, Y. et al. Optimizing neuromelanin contrast in the substantia nigra and locus coeruleus using a magnetization transfer contrast prepared 3D gradient recalled echo sequence. NeuroImage 218, 116935 (2020).

15. Turesky, T. K., Vanderauwera, J. & Gaab, N. Imaging the rapidly developing brain: Current challenges for MRI studies in the first five years of life. Dev. Cogn. Neurosci. 47, 100893 (2021).

16. Priovoulos, N. et al. Unraveling the contributions to the neuromelanin-MRI contrast. Brain Struct. Funct. 225, 2757–2774 (2020).

17. Horga, G., Wengler, K. & Cassidy, C. M. Neuromelanin-Sensitive Magnetic Resonance Imaging as a Proxy Marker for Catecholamine Function in Psychiatry. JAMA Psychiatry 78, 788–789 (2021).

18. Wengler, K. et al. Cross-Scanner Harmonization of Neuromelanin-Sensitive MRI for Multisite Studies. J. Magn. Reson. Imaging JMRI 54, 1189–1199 (2021).

19. Xing, Y., Sapuan, A., Dineen, R. A. & Auer, D. P. Life span pigmentation changes of the substantia nigra detected by neuromelanin-sensitive MRI. Mov. Disord. Off. J. Mov. Disord. Soc. 33, 1792–1799 (2018).

20. Vila, M. Neuromelanin, aging, and neuronal vulnerability in Parkinson’s disease. Mov. Disord. 34, 1440–1451 (2019).

21. Zucca, F. A. et al. Neuromelanin organelles are specialized autolysosomes that accumulate undegraded proteins and lipids in aging human brain and are likely involved in Parkinson’s disease. Npj Park. Dis. 4, 1–23 (2018).

22. Mahul-Mellier, A.-L. et al. The process of Lewy body formation, rather than simply α-synuclein fibrillization, is one of the major drivers of neurodegeneration. Proc. Natl. Acad. Sci. 117, 4971–4982 (2020).

23. Carballo-Carbajal, I. et al. Brain tyrosinase overexpression implicates age-dependent neuromelanin production in Parkinson’s disease pathogenesis. Nat. Commun. 10, 973 (2019).

24. Sasaki, M. et al. Neuromelanin magnetic resonance imaging of locus ceruleus and substantia nigra in Parkinson’s disease. Neuroreport 17, 1215–1218 (2006).

25. Shibata, E. et al. Age-related changes in locus ceruleus on neuromelanin magnetic resonance imaging at 3 Tesla. Magn. Reson. Med. Sci. MRMS Off. J. Jpn. Soc. Magn. Reson. Med. 5, 197–200 (2006).

26. Liu, K. Y. et al. In vivo visualization of age-related differences in the locus coeruleus. Neurobiol. Aging 74, 101–111 (2019).

27. Zucca, F. A. et al. Neuromelanin of the human substantia nigra: an update. Neurotox. Res. 25, 13–23 (2014).

28. Wieland, L., Fromm, S., Hetzer, S., Schlagenhauf, F. & Kaminski, J. Neuromelanin-Sensitive Magnetic Resonance Imaging in Schizophrenia: A Meta-Analysis of Case-Control Studies. Front. Psychiatry 12, (2021).

29. Therriault, J. et al. Association of Apolipoprotein E ε4 With Medial Temporal Tau Independent of Amyloid-β. JAMA Neurol. 77, 470–479 (2020).

30. Wengler, K. et al. Association between neuromelanin-sensitive MRI signal and psychomotor slowing in late-life depression. Neuropsychopharmacol. Off. Publ. Am. Coll. Neuropsychopharmacol. 46, 1233–1239 (2021).

31. Cassidy, C. M. et al. Association of locus coeruleus integrity with Braak stage and neuropsychiatric symptom severity in Alzheimer’s disease. Neuropsychopharmacology 47, 1128–1136 (2022).

