## Supplementary material for "Normative values of neuromelanin-sensitive MRI signal in older adults obtained using a standard protocol for acquisition and analysis": Table S1

| age | sex (1=male) | SN signal L | SN signal R | SN volume L | SN volume R | LC signal L | LC signal R | LC volume L | LC volume R |
|---|---|---|---|---|---|---|---|---|---|
| 86.5 | 1 | 10.09224215 | 9.207363605 | 0.584 | 0.4457 | 16.84819564 | 17.72779681 | 0.004726563 | 0.004726563 |
| 84.3 | 0 | 10.26244307 | 9.678425694 | 0.6215 | 0.5464 | 18.823511 | 13.81862126 | 0.007089844 | 0.004726563 |
| 79.4 | 0 | 10.97436705 | 9.220911503 | 0.6061 | 0.4956 | 27.55781218 | 24.81968366 | 0.007089844 | 0.005908203 |
| 78.6 | 0 | 12.12480757 | 11.01503107 | 0.636666667 | 0.561555556 | 25.4192275 | 16.43550114 | 0.007089844 | 0.007089844 |
| 75.4 | 0 | 11.95199232 | 12.25870638 | 0.6614 | 0.6212 | 30.11946778 | 24.97129656 | 0.007089844 | 0.007089844 |
| 70.7 | 0 | 9.760323811 | 10.01593103 | 0.5908 | 0.5118 | 34.12551512 | 30.16425103 | 0.007089844 | 0.007089844 |
| 71.9 | 0 | 11.81438074 | 11.38253984 | 0.6658 | 0.6016 | 25.21529354 | 11.64615344 | 0.005908203 | 0.002363281 |
| 78.5 | 1 | 10.63561859 | 11.99705038 | 0.6246 | 0.6047 | 30.72144896 | 30.74052423 | 0.007089844 | 0.007089844 |
| 53.6 | 0 | 7.565791416 | 7.375617599 | 0.4634 | 0.4186 | 26.79310411 | 13.73856198 | 0.007089844 | 0.004135742 |
| 72.3 | 1 | 10.46951113 | 11.05370646 | 0.5885 | 0.575 | 25.48034421 | 25.9427534 | 0.007030762 | 0.006499023 |
| 66.1 | 0 | 8.776576614 | 10.97381716 | 0.5365 | 0.5594 | 21.97685326 | 17.61958137 | 0.007089844 | 0.007089844 |
| 71.6 | 0 | 10.05818558 | 11.75144825 | 0.6168 | 0.646 | 17.50149116 | 13.12922637 | 0.002363281 | 0.004726563 |
| 68.5 | 0 | 12.89256611 | 12.89628925 | 0.7466 | 0.6682 | 36.91249117 | 32.16447125 | 0.007089844 | 0.007089844 |
| 81.8 | 0 | 7.854370165 | 8.994969559 | 0.4716 | 0.4962 | 21.44029148 | 19.50040599 | 0.005908203 | 0.006302083 |
| 67.4 | 0 | 10.80510798 | 11.32436504 | 0.6067 | 0.5947 | 33.39162022 | 27.00400283 | 0.007089844 | 0.007089844 |
| 68.2 | 1 | 9.297934246 | 7.891113043 | 0.5741 | 0.4215 | 27.78217077 | 19.56425428 | 0.007089844 | 0.007089844 |
| 73.4 | 0 | 7.564412866 | 10.9878728 | 0.408714286 | 0.562714286 | 19.36623752 | 19.35319752 | 0.005908203 | 0.007089844 |
| 70.1 | 0 | 8.242634201 | 8.840037537 | 0.5002 | 0.4878 | 19.74753261 | 16.22829869 | 0.007089844 | 0.005514323 |
| 75.9 | 0 | 9.75636034 | 9.813615799 | 0.5502 | 0.492 | 25.1617679 | 16.64377138 | 0.007089844 | 0.007089844 |
| 81 | 0 | 12.30079813 | 12.58494558 | 0.6503 | 0.6547 | 20.80094273 | 15.37046306 | 0.005277995 | 0.007089844 |
| 77.5 | 0 | 11.66556005 | 10.86798334 | 0.6991 | 0.5531 | 29.30628896 | 19.0128243 | 0.007089844 | 0.007089844 |
| 74.9 | 0 | 10.76434603 | 10.67324467 | 0.603 | 0.5535 | 26.23149216 | 20.69824715 | 0.007089844 | 0.007089844 |
| 61.9 | 1 | 9.666221142 | 7.309168863 | 0.5755 | 0.4272 | 16.41336445 | 10.69455598 | 0.003544922 | 0.001181641 |
| 82.8 | 0 | 9.844334793 | 8.754501915 | 0.578 | 0.4837 | 24.02118877 | 20.42472251 | 0.007089844 | 0.007089844 |
| 73.3 | 0 | 9.423601627 | 9.662572098 | 0.5484 | 0.5118 | 21.59334287 | 16.29905393 | 0.005908203 | 0.005514323 |
| 74.4 | 0 | 11.97056452 | 12.61149428 | 0.692111111 | 0.701444444 | 21.29507705 | 17.80040253 | 0.006127025 | 0.005514323 |
| 69 | 0 | 8.925805759 | 8.130561829 | 0.514 | 0.4341 | 24.70275432 | 19.92064178 | 0.007089844 | 0.007089844 |
| 71.6 | 0 | 9.09779768 | 10.4973237 | 0.5358 | 0.5869 | 22.4028255 | 17.45184027 | 0.007089844 | 0.005908203 |
| 72 | 0 | 10.1090642 | 10.33043823 | 0.5905 | 0.5639 | 17.78703742 | 15.81657678 | 0.005908203 | 0.006223307 |
| 68.1 | 1 | 10.26231899 | 9.465454578 | 0.5909 | 0.5249 | 29.17473018 | 22.33720129 | 0.007089844 | 0.006302083 |
| 75.6 | 0 | 11.76258135 | 12.38584251 | 0.7104 | 0.6411 | 19.42047708 | 14.86820124 | 0.006026367 | 0.005041667 |
| 71 | 0 | 9.681323528 | 10.74522924 | 0.5635 | 0.5678 | 24.74528812 | 22.03905478 | 0.007089844 | 0.007089844 |
| 75.5 | 1 | 11.32888498 | 10.52710199 | 0.6205 | 0.5329 | 20.70175305 | 17.36041116 | 0.007089844 | 0.005514323 |
| 85.1 | 0 | 11.65929174 | 9.144820499 | 0.6417 | 0.5158 | 24.26200901 | 21.78051859 | 0.006538411 | 0.007089844 |
| 65.8 | 0 | 9.486903826 | 10.75797346 | 0.559333333 | 0.590444444 | 24.38726701 | 24.37261062 | 0.007089844 | 0.007089844 |
| 73.1 | 0 | 11.03274565 | 11.74041853 | 0.6506 | 0.6236 | 19.29047167 | 15.17901123 | 0.006302083 | 0.007089844 |
| 79.4 | 0 | 8.479627323 | 9.145263958 | 0.4706 | 0.4576 | 18.53140729 | 10.16968708 | 0.003938802 | 0.001181641 |
| 67.5 | 1 | 11.35213003 | 11.36882219 | 0.6612 | 0.6019 | 21.91283325 | 22.87421917 | 0.007089844 | 0.007089844 |
| 65.8 | 0 | 11.58098755 | 11.09259348 | 0.6867 | 0.5658 | 28.29470634 | 29.43815514 | 0.007089844 | 0.007089844 |
| 68.8 | 1 | 8.211422825 | 8.902625179 | 0.4684 | 0.4738 | 25.38365766 | 20.83036085 | 0.007089844 | 0.007089844 |
| 72 | 1 | 7.476081276 | 7.035644341 | 0.4255 | 0.3974 | 22.95774167 | 16.11599262 | 0.006302083 | 0.005514323 |
| 68.3 | 0 | 8.850000668 | 8.802823162 | 0.51 | 0.452 | 23.80277147 | 24.1851896 | 0.007089844 | 0.007089844 |
| 67.8 | 0 | 9.097699642 | 10.203092 | 0.5207 | 0.532 | 21.12464155 | 17.6979804 | 0.005514323 | 0.007089844 |
| 69.6 | 0 | 8.487627602 | 9.339304638 | 0.4861 | 0.4353 | 17.62470543 | 18.00213203 | 0.003151042 | 0.005514323 |
| 68.1 | 1 | 9.874359608 | 9.936196136 | 0.5715 | 0.5498 | 23.16855997 | 26.88827246 | 0.007011068 | 0.007089844 |
| 65.7 | 1 | 11.71396809 | 11.40178833 | 0.6399 | 0.5736 | 29.1631037 | 28.87162 | 0.007089844 | 0.007089844 |
| 72.3 | 0 | 12.84026785 | 13.4689539 | 0.7143 | 0.7041 | 23.68524676 | 21.30232602 | 0.006380859 | 0.007089844 |
| 65.2 | 0 | 10.22940416 | 10.17465124 | 0.6482 | 0.5999 | 27.33065456 | 27.87652493 | 0.007089844 | 0.007089844 |
| 86.5 | 0 | 11.87911715 | 10.92600373 | 0.672857143 | 0.601428571 | 21.92833757 | 16.9436564 | 0.006302083 | 0.006302083 |
| 78.5 | 0 | 9.806978226 | 9.911590576 | 0.5674 | 0.5296 | 30.54719403 | 21.76632755 | 0.007089844 | 0.007089844 |
| 76.1 | 0 | 7.44078021 | 7.097746754 | 0.4342 | 0.3732 | 19.23105411 | 14.52239692 | 0.005908203 | 0.007089844 |
| 71.4 | 0 | 11.15329666 | 10.87509804 | 0.6534 | 0.5743 | 20.0258183 | 19.74810849 | 0.007089844 | 0.006302083 |
| 74.9 | 1 | 9.512755489 | 10.00515785 | 0.5271 | 0.5088 | 17.74420299 | 17.56801553 | 0.005081055 | 0.004726563 |
| 70.3 | 1 | 10.56963472 | 11.12330275 | 0.5729 | 0.5716 | 38.51014674 | 30.17367795 | 0.007089844 | 0.007089844 |
| 57.2 | 0 | 11.42995663 | 10.52130957 | 0.7035 | 0.5796 | 23.20528229 | 17.53076747 | 0.006302083 | 0.005908203 |
| 66 | 0 | 9.926638508 | 10.1702611 | 0.5716 | 0.5079 | 19.15397008 | 20.96032058 | 0.007089844 | 0.007089844 |
| 72.5 | 0 | 9.901811886 | 11.21640034 | 0.6403 | 0.601 | 23.46029351 | 15.38394456 | 0.005514323 | 0.004726563 |
| 76.5 | 0 | 9.793280029 | 10.73396111 | 0.5606 | 0.574 | 27.96632044 | 23.60629285 | 0.007089844 | 0.007089844 |
| 75.2 | 1 | 11.40380116 | 9.268073559 | 0.6411 | 0.4661 | 24.90703911 | 16.92960039 | 0.007089844 | 0.003544922 |
| 77.6 | 0 | 11.4557333 | 11.55919895 | 0.6867 | 0.6057 | 29.90466729 | 18.82540405 | 0.007089844 | 0.007089844 |
| 66.7 | 1 | 10.4109787 | 11.46955395 | 0.5741 | 0.5626 | 28.59161059 | 29.16175291 | 0.007089844 | 0.007089844 |
| 65.7 | 0 | 11.73358955 | 12.6968667 | 0.6704 | 0.6131 | 23.21255468 | 19.18205296 | 0.007089844 | 0.007089844 |
| 79.4 | 0 | 10.14914827 | 8.973592949 | 0.5995 | 0.5239 | 21.15494092 | 19.27195599 | 0.007089844 | 0.005986979 |
| 65.6 | 0 | 11.85486526 | 10.74967365 | 0.6198 | 0.5837 | 28.45641797 | 19.62209518 | 0.007089844 | 0.006302083 |
| 66 | 1 | 11.72167416 | 12.19568615 | 0.6709 | 0.6192 | 17.55542417 | 14.16550294 | 0.003938802 | 0.005514323 |
| 67.3 | 1 | 9.815086079 | 10.29347782 | 0.5618 | 0.533 | 27.28601813 | 25.45671135 | 0.007089844 | 0.007089844 |
| 68.3 | 0 | 9.666300201 | 11.46186771 | 0.547 | 0.5961 | 25.41553944 | 20.63664205 | 0.007089844 | 0.007089844 |
| 65.1 | 0 | 9.648914814 | 9.006492519 | 0.5853 | 0.4823 | 32.11793981 | 24.3839774 | 0.007089844 | 0.007089844 |
| 68.7 | 1 | 7.154340982 | 8.651348782 | 0.4083 | 0.4346 | 27.83445666 | 27.17376053 | 0.007089844 | 0.007089844 |
| 67 | 0 | 10.31777353 | 10.52669697 | 0.6432 | 0.565 | 22.88437409 | 18.31196658 | 0.00437207 | 0.003544922 |
| 70.5 | 0 | 12.29263487 | 12.52867575 | 0.6959 | 0.6288 | 21.70878112 | 20.57009627 | 0.007089844 | 0.007089844 |

| | | | | | | | | | |
|---|---|---|---|---|---|---|---|---|---|
| 68.3 | 0 | 10.92767448 | 11.15219297 | 0.6964 | 0.6225 | 25.07063583 | 22.50657268 | 0.007089844 | 0.007089844 |
| 72.1 | 1 | 6.262666273 | 8.138277721 | 0.3643 | 0.4698 | 25.54553757 | 18.29219004 | 0.006302083 | 0.006302083 |
| 73.3 | 0 | 8.351290894 | 10.08846006 | 0.4629 | 0.5307 | 26.38701876 | 17.19153419 | 0.007089844 | 0.004844727 |
| 68.6 | 0 | 9.394157696 | 11.7412385 | 0.5324 | 0.5954 | 20.11841238 | 15.07752985 | 0.004844727 | 0.007089844 |
| 74.4 | 0 | 11.72980137 | 10.8571641 | 0.6963 | 0.5834 | 30.79860106 | 25.25053725 | 0.007089844 | 0.007089844 |
| 72.6 | 1 | 12.15648909 | 12.28369932 | 0.6795 | 0.635 | 26.10587887 | 21.32613242 | 0.007089844 | 0.007089844 |
| 72.9 | 0 | 12.41839542 | 12.95223627 | 0.652 | 0.6357 | 28.89069945 | 22.2492217 | 0.005671875 | 0.007089844 |
| 71.5 | 1 | 9.247956181 | 10.88604231 | 0.5224 | 0.5469 | 27.76715298 | 26.12362007 | 0.007089844 | 0.007089844 |
| 81.9 | 0 | 10.29779301 | 10.20762014 | 0.5605 | 0.5268 | 33.82951453 | 26.71231568 | 0.007089844 | 0.007089844 |
| 79.9 | 1 | 9.237420845 | 9.755372143 | 0.5294 | 0.5394 | 35.53542763 | 29.32264954 | 0.007089844 | 0.007089844 |
| 70.2 | 0 | 10.69444942 | 9.492100716 | 0.6263 | 0.5096 | 24.28757151 | 23.14681709 | 0.007089844 | 0.007089844 |
| 67.2 | 0 | 9.719609451 | 11.35179939 | 0.5244 | 0.545 | 24.2691768 | 21.23383348 | 0.004726563 | 0.004805339 |
| 74.1 | 0 | 9.720587158 | 10.01206427 | 0.5742 | 0.5487 | 19.50717906 | 13.30682375 | 0.003938802 | 0.002363281 |
| 75 | 1 | 10.79288549 | 11.48334541 | 0.6062 | 0.5893 | 27.470044 | 14.46397218 | 0.007089844 | 0.007089844 |
| 66.8 | 0 | 12.7463068 | 12.3106142 | 0.7438 | 0.6794 | 21.45452745 | 14.97203484 | 0.007089844 | 0.004726563 |
| 73.2 | 0 | 9.550484848 | 11.94797468 | 0.5456 | 0.5785 | 21.544495 | 14.42203477 | 0.003544922 | 0.005908203 |
| 76.8 | 0 | 9.759734249 | 10.35079556 | 0.5773 | 0.562 | 33.05856749 | 31.88565701 | 0.007089844 | 0.007089844 |
| 66.8 | 1 | 10.0810092 | 10.92838106 | 0.5929 | 0.6021 | 24.47314873 | 16.43236841 | 0.007089844 | 0.006302083 |
| 71.3 | 1 | 10.63707418 | 12.25516243 | 0.6114 | 0.6062 | 24.60179682 | 21.69984125 | 0.007089844 | 0.007089844 |
| 68.5 | 1 | 11.25606117 | 8.727684212 | 0.6418 | 0.4761 | 24.56875503 | 19.55932677 | 0.007089844 | 0.006538411 |
| 70.2 | 1 | 6.002875423 | 6.399903393 | 0.3545 | 0.3299 | 22.98566706 | 21.48311622 | 0.007089844 | 0.007089844 |
| 66.9 | 1 | 8.996471214 | 8.357442093 | 0.5283 | 0.4503 | 21.33115619 | 17.39023348 | 0.004096354 | 0.007089844 |
| 67.8 | 0 | 10.79203091 | 11.99423351 | 0.6077 | 0.5906 | 24.35293801 | 18.16749369 | 0.007089844 | 0.007089844 |
| 66.7 | 1 | 9.265492153 | 9.754935837 | 0.551 | 0.5513 | 19.00043033 | 19.49371561 | 0.004726563 | 0.005514323 |
| 73.2 | 0 | 13.42492685 | 11.67453146 | 0.7324 | 0.5861 | 29.05696556 | 27.0543618 | 0.007089844 | 0.007089844 |
| 73.6 | 1 | 6.600211859 | 6.587846375 | 0.4 | 0.3402 | 27.75450051 | 22.09380942 | 0.007089844 | 0.006538411 |
| 65.3 | 1 | 10.70785866 | 10.20743465 | 0.6061 | 0.5393 | 28.89327824 | 23.95235563 | 0.007089844 | 0.007089844 |
| 73.7 | 0 | 10.21944523 | 9.916686249 | 0.6131 | 0.5281 | 21.25516973 | 18.53945389 | 0.007089844 | 0.005908203 |
| 78.6 | 0 | 12.82620687 | 12.10999899 | 0.7039 | 0.5713 | 26.49773374 | 19.72774367 | 0.005514323 | 0.005514323 |
| 60.8 | 1 | 10.16960812 | 11.9040555 | 0.5805 | 0.6067 | 28.18661769 | 18.98897948 | 0.007089844 | 0.005671875 |
| 71.5 | 0 | 8.911065292 | 9.768029118 | 0.5181 | 0.5238 | 23.30997884 | 19.54131559 | 0.004726563 | 0.005908203 |
| 63.4 | 0 | 10.3343667 | 10.11013765 | 0.6481 | 0.554 | 14.31481446 | 13.20895031 | 0.001654297 | 0.003938802 |
| 73.6 | 0 | 7.770002937 | 8.387005615 | 0.4453 | 0.4236 | 17.0167252 | 17.39643726 | 0.003544922 | 0.006144531 |
| 66.6 | 0 | 9.737930298 | 9.530849743 | 0.5967 | 0.5187 | 20.19804701 | 20.7014776 | 0.003544922 | 0.005908203 |
| 74.3 | 0 | 8.963054466 | 8.355564976 | 0.491 | 0.4668 | 31.80753395 | 23.12892149 | 0.007089844 | 0.007089844 |
| 67.4 | 0 | 8.059640789 | 8.779325962 | 0.4753 | 0.4936 | 21.86176946 | 16.27651766 | 0.006302083 | 0.007089844 |
| 67.1 | 0 | 10.85876417 | 10.58497829 | 0.6367 | 0.549 | 21.59933984 | 21.80849254 | 0.007089844 | 0.006302083 |
| 77.5 | 0 | 9.405270863 | 9.765023518 | 0.5748 | 0.5394 | 24.14058998 | 19.89144961 | 0.007089844 | 0.007089844 |
| 55.5 | 0 | 11.27320099 | 12.0523798 | 0.6515 | 0.6235 | 27.73605516 | 22.39925325 | 0.007089844 | 0.007089844 |
| 62.7 | 0 | 9.2414258 | 10.84516973 | 0.5814 | 0.6225 | 20.1254575 | 15.50440557 | 0.007089844 | 0.005514323 |
| 66 | 0 | 11.09198465 | 11.92393541 | 0.6253 | 0.6162 | 25.29309168 | 20.85108583 | 0.007089844 | 0.007089844 |
| 71.8 | 1 | 8.175478935 | 8.664843877 | 0.474222222 | 0.456888889 | 29.61669686 | 24.94563725 | 0.007089844 | 0.007089844 |
| 70.9 | 0 | 9.578463364 | 9.871462536 | 0.5505 | 0.5124 | 26.78288564 | 25.08625343 | 0.007089844 | 0.007089844 |
| 67.5 | 0 | 10.07033978 | 11.84148912 | 0.5753 | 0.6107 | 16.5275763 | 16.5275763 | 0.002363281 | 0.007089844 |
| 71.1 | 0 | 10.39427843 | 9.077067757 | 0.5701 | 0.4513 | 17.07210444 | 17.08105907 | 0.004726563 | 0.007089844 |
| 66.3 | 1 | 10.19977627 | 10.33298283 | 0.5734 | 0.5216 | 31.65910438 | 22.7045315 | 0.007089844 | 0.007089844 |
| 67.5 | 0 | 5.814394772 | 6.41652149 | 0.331125 | 0.322125 | 24.47054889 | 16.41471883 | 0.007089844 | 0.005908203 |
| 68.1 | 0 | 8.73984623 | 9.709532642 | 0.5126 | 0.5092 | 20.24594255 | 14.83244918 | 0.005908203 | 0.005908203 |
| 71.8 | 1 | 9.447615351 | 8.647656032 | 0.532 | 0.463857143 | 28.49645423 | 23.34937091 | 0.005908203 | 0.007089844 |
| 75.9 | 0 | 10.2957428 | 11.90905695 | 0.5756 | 0.6219 | 17.62100627 | 13.65281922 | 0.003938802 | 0.006459635 |
| 78.8 | 1 | 8.069597138 | 6.69241635 | 0.468 | 0.362555556 | 21.30512457 | 21.97470488 | 0.005514323 | 0.005339265 |
| 65.1 | 0 | 11.76263018 | 12.35786142 | 0.6849 | 0.6448 | 18.66802506 | 18.56869757 | 0.004805339 | 0.004844727 |
| 80.7 | 0 | 8.823777485 | 10.92113533 | 0.495 | 0.5586 | 23.22677801 | 13.32237865 | 0.003938802 | 0.004726563 |
| 71.9 | 1 | 9.292297363 | 7.815124512 | 0.533666667 | 0.432111111 | 29.8663231 | 24.36298965 | 0.007089844 | 0.007089844 |
| 76.2 | 1 | 8.978890038 | 8.559309196 | 0.5197 | 0.4752 | 21.25672691 | 13.66445623 | 0.005908203 | 0.005908203 |
| 79.1 | 1 | 8.610056686 | 9.825812817 | 0.5022 | 0.5225 | 20.3653502 | 16.03819062 | 0.005514323 | 0.006302083 |
| 69 | 0 | 9.184625244 | 10.81545391 | 0.5072 | 0.578 | 19.81427744 | 13.95501494 | 0.007089844 | 0.004726563 |
| 70.3 | 1 | 10.25763988 | 11.58365965 | 0.5976 | 0.5966 | 29.25663427 | 23.95419158 | 0.007089844 | 0.007089844 |
| 69.1 | 0 | 10.34258089 | 12.41884136 | 0.5489 | 0.6172 | 25.29650857 | 19.59654907 | 0.006302083 | 0.006302083 |
| 69.9 | 0 | 13.20075274 | 12.5230835 | 0.7324 | 0.6394 | 36.17095242 | 28.71807774 | 0.007089844 | 0.007089844 |
| 80.5 | 1 | 7.512774876 | 7.98052597 | 0.44 | 0.428428571 | 32.58913174 | 32.12133206 | 0.007089844 | 0.007089844 |
| 78.6 | 0 | 8.477090168 | 9.007842636 | 0.4836 | 0.4875 | 17.0991046 | 16.53601803 | 0.005908203 | 0.004253906 |
| 71.9 | 0 | 8.157921696 | 9.796482182 | 0.4745 | 0.4941 | 26.13571127 | 20.28043643 | 0.007089844 | 0.007089844 |
| 64.4 | 0 | 10.91474329 | 11.04024683 | 0.598 | 0.559428571 | 25.49652383 | 22.9951481 | 0.007089844 | 0.007089844 |
| 67.1 | 0 | 12.17432852 | 12.11823854 | 0.7305 | 0.6582 | 28.32405657 | 24.44069818 | 0.005908203 | 0.007089844 |
| 65.7 | 1 | 8.600915432 | 9.458952522 | 0.5327 | 0.5146 | 19.81992637 | 17.63810987 | 0.004726563 | 0.005277995 |
| 75 | 1 | 8.66205101 | 8.1564188 | 0.5044 | 0.4085 | 21.25297904 | 16.99647377 | 0.006302083 | 0.006302083 |
| 65.7 | 0 | 10.0866765 | 8.980148888 | 0.5712 | 0.4556 | 27.58323818 | 18.39653082 | 0.005908203 | 0.005908203 |
| 71.8 | 0 | 8.978473568 | 9.598110008 | 0.5034 | 0.4751 | 25.85324891 | 21.30228817 | 0.007089844 | 0.007089844 |
| 62.3 | 0 | 9.748620415 | 10.01632738 | 0.5226 | 0.5056 | 25.34983565 | 19.56137046 | 0.006302083 | 0.006302083 |
| 66.2 | 1 | 10.64649086 | 11.99114628 | 0.6028 | 0.5932 | 21.85961887 | 15.97173062 | 0.005514323 | 0.007089844 |
| 59.7 | 1 | 10.5722003 | 11.68542223 | 0.643 | 0.6476 | 30.22885203 | 29.62601826 | 0.007089844 | 0.007089844 |

| | | | | | | | | | |
|---|---|---|---|---|---|---|---|---|---|
| 68.7 | 0 | 9.546656895 | 9.824629402 | 0.5301 | 0.5449 | 17.07965672 | 13.60553309 | 0.003544922 | 0.003544922 |
| 73.2 | 1 | 8.536703968 | 8.232372379 | 0.4831 | 0.419 | 25.52978575 | 33.00245404 | 0.007089844 | 0.007089844 |
| 71.5 | 1 | 10.38927193 | 10.49403486 | 0.5885 | 0.5511 | 37.83468907 | 31.83581755 | 0.004726563 | 0.004096354 |
| 81.4 | 0 | 10.61729679 | 11.29257622 | 0.603 | 0.594 | 28.64352465 | 23.19462275 | 0.007089844 | 0.007089844 |
| 70.5 | 1 | 10.17783006 | 8.622163879 | 0.577666667 | 0.457111111 | 26.65044044 | 20.67243084 | 0.007089844 | 0.007089844 |
| 59.9 | 0 | 11.55121717 | 11.01920834 | 0.6426 | 0.5814 | 27.31228411 | 24.70243871 | 0.007089844 | 0.007089844 |
| 80.1 | 0 | 9.821352768 | 10.32024336 | 0.5529 | 0.5699 | 30.43453082 | 20.70568338 | 0.007089844 | 0.007089844 |
| 76.2 | 0 | 11.32036915 | 11.14948931 | 0.6441 | 0.5454 | 31.38389215 | 24.32636507 | 0.007089844 | 0.007089844 |
| 70.7 | 0 | 8.596563435 | 10.7824398 | 0.4497 | 0.5327 | 23.50184351 | 16.08444355 | 0.007089844 | 0.007089844 |
